# Resistance Training Diminishes Mitochondrial Adaptations to Subsequent Endurance Training

**DOI:** 10.1101/2023.04.06.535919

**Authors:** Paulo H. C. Mesquita, Joshua S. Godwin, Bradley A. Ruple, Casey L. Sexton, Mason C. McIntosh, Breanna J. Mueller, Shelby C. Osburn, C. Brooks Mobley, Cleiton A. Libardi, Kaelin C. Young, L. Bruce Gladden, Michael D. Roberts, Andreas N. Kavazis

## Abstract

We investigated the effects of performing a period of resistance training (RT) on the performance and molecular adaptations to a subsequent period of endurance training (ET). Twenty-five young adults were divided into RT+ET (n=13), which underwent seven weeks of RT followed by seven weeks of ET, and ET-only (n=12), which performed seven weeks of ET. Body composition, endurance performance, and muscle biopsies were collected before RT (T1, baseline for RT+ET), before ET (T2, post RT for RT+ET and baseline for ET), and after ET (T3). Immunohistochemistry was performed to determine fiber cross-sectional area (fCSA), myonuclear content, myonuclear domain size, satellite cell number, and mitochondrial content. Western blots were used to quantify markers of mitochondrial remodeling. Citrate synthase activity and markers of ribosome content were also investigated. Resistance training improved body composition and strength, increased vastus lateralis thickness, mixed and type II fCSA, myonuclear number, markers of ribosome content, and satellite cell content (p<0.050). In response to ET, both groups similarly decreased body fat percentage and improved endurance performance (e.g., VO_2_max, and speed at which the onset of blood lactate accumulation occurred during the VO_2_max test). Levels of mitochondrial complexes I-IV in the ET-only group increased 32-66%, while the RT+ET group increased 1-11%. Additionally, mixed fiber relative mitochondrial content increased 15% in the ET-only group but decreased 13% in the RT+ET group. In conclusion, RT performed prior to ET had no additional benefits to ET adaptations. Moreover, prior RT seemed to impair mitochondrial adaptations to ET.

**KEY POINTS SUMMARY:**

- Resistance training is largely underappreciated as a method to improve endurance performance, despite reports showing it may improve mitochondrial function.
- Although several concurrent training studies are available, in this study we investigated the effects of performing a period resistance training on the performance and molecular adaptations to subsequent endurance training.
- Prior resistance training did not improve endurance performance and impaired most mitochondrial adaptations to subsequent endurance training, but that seemed to be a result of detraining from resistance training.

## INTRODUCTION

Endurance performance is determined by a complex interaction of physiological, biomechanical, and neuromuscular factors. Maximal oxygen consumption (VO_2_max), lactate threshold, and running economy are widely considered the main limiting factors of endurance performance (1, 2). Skeletal muscle oxidative phosphorylation capacity, which in turn is determined by mitochondrial volume density and function, is also considered a strong predictor of endurance performance (3).

A variety of training paradigms may be used to improve endurance performance including moderate intensity continuous training (MICT) and high-intensity interval training (HIIT) (4-6). Resistance training (RT), on the other hand, has long been underappreciated in regard to enhancing endurance performance and is often not a part of the training program of high-level athletes (7). However, several studies have shown a beneficial effect of RT on endurance performance, which is usually linked to an improvement of running economy through neuromuscular adaptations (8-10).

The enhancement of endurance performance through RT may affect attributes other than running economy. Different studies have shown that RT may also lead to positive mitochondrial adaptations (11-14). However, endurance performance was not investigated in these studies. Interestingly, a study conducted by Lee et al. (15) in rats found that RT promoted enhanced mitochondrial adaptations to a subsequent block of RT, which seemed to be related to increased myonuclear number per myofiber achieved in the first block of training. While these data are promising, it is currently unknown whether RT enhances mitochondrial adaptations to subsequent endurance training (ET) in humans and whether the enhanced mitochondrial adaptations would lead to better endurance performance. In addition, while several studies have investigated the effects of concurrent training, when both RT and ET are combined within the same training session or program, no study to date has employed a design that investigated RT-only followed by ET-only. This is especially relevant considering a recent study reported that prior ET facilitated adaptations to a subsequent period of RT (16).

Therefore, the purpose of this study was to investigate the effects of prior RT on the molecular and performance adaptations to subsequent ET in humans. We hypothesized that RT prior to ET would augment skeletal muscle mitochondrial adaptations to ET, ultimately leading to improved endurance performance.

## MATERIALS AND METHODS

### Ethical approval

The current study was reviewed and approved by the Institutional Review Board at Auburn University (Protocol # 21-390 FB) and conformed to the standards of the Declarations of Helsinki, except that it was not registered as a clinical trial.

### Participants

Twenty-five healthy young male participants (baseline characteristics in Table 1) were recruited to participate in this study. Participants should not have participated in structured (more than once weekly for at least two months) RT over the last three years or ET over the last six months prior to joining the current study. All participants were informed of the procedures and risks of the current study before providing written consent.

**Table 1.** Participant characteristics obtained during the familiarization session.

| Characteristic | Overall (n=25) | RT+ET (n=13) | ET-only (n=12) |
| --- | --- | --- | --- |
| Age (years) | 23 ± 4 | 23 ± 4 | 24 ± 4 |
| Body mass (kg) | 84.8 ± 18.0 | 81.2 ± 16.8 | 88.6 ± 19.2 |
| Height (cm) | 181 ± 8 | 180 ± 9 | 181 ± 9 |
| BMI (kg/m<sup>2</sup>) | 25.9 ± 4.6 | 25.2 ± 5 | 26.7 ± 4.2 |
| VO<sub>2</sub>peak (ml/kg/min) | 39.7 ± 7.6 | 40.4 ± 8.9 | 38.9 ± 6.2 |

Abbreviations: RT+ET, group that performed 7 weeks of resistance training followed by 7 weeks of endurance training; ET-only, group that performed 7 weeks of endurance training only; BMI, body mass index; VO<sub>2</sub>peak, peak aerobic capacity

### Familiarization session

Participants visited the laboratory to become familiarized with the exercises and tests used in the study. First, participants performed a maximal cardiorespiratory test on a motorized treadmill. The incremental treadmill test was composed of several two-minute stages and started with a fast-paced walk (6.4 km/h, 0% inclination) as a warm-up for three minutes. After that, the speed of the treadmill increased by 1 km/h and inclination by 1% after every stage until the participant reached volitional exhaustion. There was a 30-second break after each stage, during which participants stopped running and reported their ratings of perceived exertion (RPE) according to Borg’s CR10 scale (17). Peak oxygen consumption (VO_2peak_) was determined by the highest 30-second average value using a metabolic cart (True Max 2400, ParvoMedics, Salt Lake City, UT, USA). After the maximal cardiorespiratory test, participants were taught how to properly perform the leg press, bench press, leg extension, cable-bar pull-down, and leg curl exercises. Participants were allowed to perform a few sets and repetitions until they demonstrated proper lifting technique.

### Experimental design

VO_2peak_ values obtained during the familiarization visit were used to assign participants to each group in a balanced manner, RT+ET (n=13, VO_2_peak = 40.4 ± 8.9 ml/kg/min) and ET-only (n=12, VO_2_peak = 38.9 ± 6.2 ml/kg/min). Importantly, there was no significant difference between groups (p=0.634).

Participants in the RT+ET group completed seven weeks of RT followed by seven weeks of ET. Participants in the ET-only group performed seven weeks of ET. During each of the three testing sessions (details provided below), participants underwent a battery of assessments in two visits. The first visit included height, body mass, full-body dual-energy X-ray absorptiometry (DEXA), ultrasound of the right vastus lateralis (VL), and a biopsy from the right VL. In the second visit, participants performed a maximal cardiorespiratory test and 3-repetition maximum strength tests. The order of the tests was the same for all time-points. The first and second visits of the testing sessions occurred within 48 to 96 hours and within one week after the last training session, respectively. The experimental design is depicted in Fig. 1.

**Figure 1.**
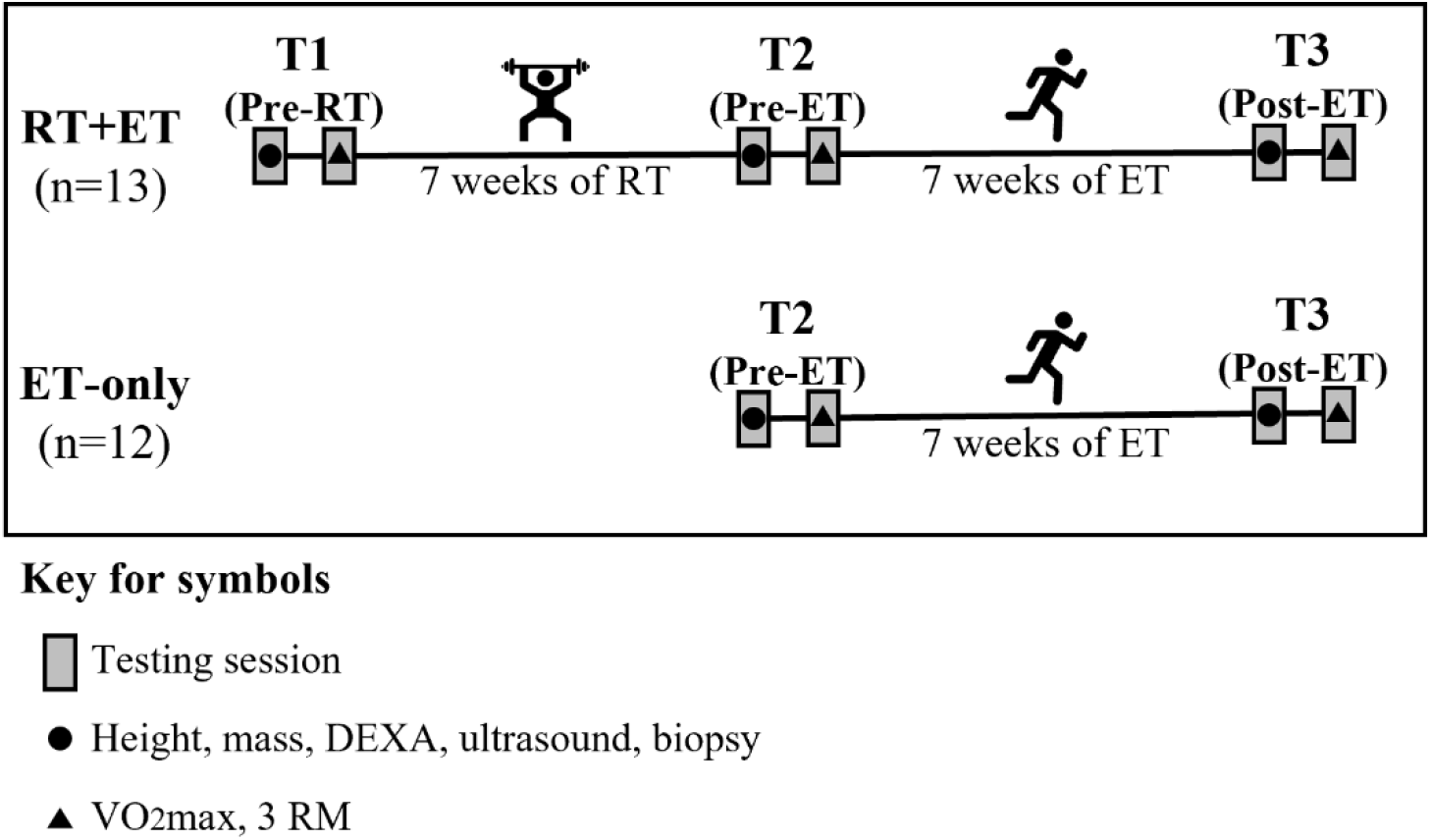
Experimental Design. Abbreviations: RT+ET, group that performed 7 weeks of resistance training followed by 7 weeks of endurance training; ET-only, group that performed 7 weeks of endurance training only; RT, Resistance Training; ET, Endurance Training *Testing sessions (T1, T2, and T3)*

Testing sessions were composed of two visits on different days. During the first day, participants reported to the laboratory following at least four hours of food deprivation. Body mass and height were assessed with a digital scale (Seca 769; Hanover, MD, USA). DEXA (Lunar Prodigy; GE Corporation, Fairfield, CT, USA) was performed to determine lean body mass, fat mass, and body fat percentage. Following the DEXA scan, real-time B-mode ultrasonography (NextGen LOGIQe R8, GE Healthcare, Chicago, IL, USA) was used to determine the thickness of the VL of the right leg as previously described by our laboratory (18, 19). Measurements were taken at the midway point between the iliac crest and proximal patella. After the ultrasound scans, skeletal muscle biopsy samples were collected from the right VL at the same location of the ultrasound imaging using a 5-gauge Bergstrom needle. Briefly, participants laid in the supine position on an athletic training table and the upper thigh was shaven and cleaned with 70% isopropanol before receiving a 0.8 mL injection of 1% lidocaine. Participants rested for 5-10 minutes for the lidocaine to take effect before the area was cleaned with chlorhexidine and a pilot incision through the dermis was made with a sterile No. 11 surgical blade (AD Surgical; Sunnyvale CA, USA). Approximately 50-100 mg of skeletal muscle tissue was collected, immediately teased of blood and connective tissue, and separated for histological and biochemical analysis. Mounting for histology in optimal cutting temperature (OCT) media occurred as previously described by our laboratory (14). A separate ∼20-40 mg tissue sample was placed in pre-labelled foil and flash-frozen in liquid nitrogen for Western blotting and biochemical analyses described below. Finally, ∼10 mg of muscle was fixed in 4% paraformaldehyde for 48 hours at room temperature for single fiber analysis and is further described below. Notably, removal of tissue and all tissue processing occurred within a 5-minute period. Furthermore, OCT and flash frozen foil samples were removed from liquid nitrogen throughout the day during muscle collections and stored at -80°C for later analyses.

During the second day of testing, participants performed a maximal cardiorespiratory test as previously described in the familiarization session. For the testing session, blood was also collected from the participants’ fingertips at rest, after completion of each stage, and at the end of the test. A handheld lactate analyzer device (Lactate Plus, Nova Biomedical) was used to obtain blood lactate concentration values. Blood lactate values were used to determine the speed and inclination corresponding to the onset of blood lactate accumulation (OBLA, i.e., 4 mmol/L) using the Lactater package in RStudio. In addition, a validation step was conducted at the end of the test. After completing the test, participants rested for 10 minutes, were connected to the metabolic cart again and ran for as long they could at a speed and inclination corresponding to the stage following the stage they stopped during the test. This step was included as a verification method to ensure that participants reached maximal oxygen consumption (VO_2_max). The highest 30-second average oxygen consumption value obtained during the test was considered the participants’ VO_2_max. After the VO_2_max test, participants completed three-repetition maximum (3RM) strength tests for the leg press, bench press, and leg extension exercises. Participants performed two sets for warm-up and had up to five trials per exercise to reach 3RM values with three to five minutes of rest between trials. Proper range of motion was assessed for each exercise during the warm-up with the aid of a measurement tape, and repetitions were considered valid if participants reached appropriate ranges of motion.

### Resistance training

Resistance training was performed twice weekly by the RT+ET group only, and each training session included leg press, bench press, leg extension, cable pull-down, and leg curls. Sets of six repetitions were performed for the exercises targeting quadriceps muscles (i.e., leg press and leg extension), while three sets of ten repetitions were performed for the other exercises (i.e., bench press, cable pull-down, leg curls). Volume and load for the quadriceps were progressively increased throughout the seven weeks and can be seen in Table 2. Although the load increment was pre-planned as shown, participant feedback was taken into consideration for load adjustments. After each set for each exercise, participants reported their repetitions in reserve (RIR) by answering how many more repetitions they think they could have done (20). If RIR > 2, the load was increased by approximately 5-10 lbs for upper-body exercises and 10-20 lbs for lower-body exercises. If participants failed to perform the programmed number of repetitions, load was decreased in a similar fashion. Participants rested for two to three minutes between sets of exercises. Appropriate range of motion was ensured using the range of motion recorded at T1-testing.

**Table 2.** Strength training volume and load progression.

| Week | Day | Sets x reps* | 1RM |
| --- | --- | --- | --- |
| 1 | 1 | 6 x 6 | 70% |
|  | 2 | 6 x 6 | 70% |
| 2 | 3 | 7 x 6 | 75% |
|  | 4 | 7 x 6 | 75% |
| 3 | 5 | 8 x 6 | 80% |
|  | 6 | 8 x 6 | 80% |
| 4 | 7 | 9 x 6 | 85% |
|  | 8 | 8 x 6 | 85% |
| 5 | 9 | 9 x 6 | 88% |
|  | 10 | 9 x 6 | 88% |
| 6 | 11 | 10 x 6 | 91% |
|  | 12 | 9 x 6 | 91% |
| 7 | 13 | 10 x 6 | 95% |
|  | 14 | 10 x 6 | 95% |

Abbreviation: 1RM, one-repetition maximum. Symbol: \*Total number of sets and reps per session for exercises targeting quadriceps (e.g., 6 x 6 = 3 sets of 6 repetitions for the leg press and 3 sets of 6 repetitions for the leg extension; 9 x 6 = 5 sets of 6 repetitions for the leg press and 4 sets of 6 repetitions for the leg extension).

### Endurance training

All participants performed seven weeks of a high-intensity interval training (HIIT)-based ET on a motorized treadmill, and in RT+ET participants, this training occurred the week immediately following their seven week-RT period. A HIIT-based training protocol was chosen because HIIT induces mitochondrial and cardiovascular adaptations in a time-efficient manner (6, 21). For each training session, participants started with a 3-minute warm-up, followed by 5-10 sets (5 sets in the 1st week; 8 sets in the 2nd week; 9 sets in the 3rd week; 10 sets for the remaining weeks) of 1 minute running at a high intensity interspersed by 1.5 to 3 minutes running at a low intensity. The intensity of the “sprints” and the recovery was determined using the speed and inclination values achieved in the VO_2_max test (Table 3). Similar to the RT program, participant feedback was taken into consideration to adjust the intensity of training. At the end of each “sprint” bout, participants rated their perceived exertion (RPE) using the CR-10 Borg Scale. In the first week, if participant’s final RPE was lower than 5 (“strong”), the intensity of the “sprint” bout was increased in the next training session by 5%. From the second week onward, if participant’s final RPE was lower than 7 (“very strong”), intensity (i.e., treadmill speed) was also increased by 5%. If participants were not able to complete the programmed number of “sprints”, intensity was decreased by 5%. The ET program can be seen in Table 3:

**Table 3.** Endurance training program.

| Week | Frequency<br>(times/week) | Sets | Effort dur<br>(min) | Effort int<br>(VO<sub>2</sub>max) | Recovery<br>dur (min) | Recovery int<br>(VO<sub>2</sub>max) |
| --- | --- | --- | --- | --- | --- | --- |
| 1 | 2 | 5 | 1 | 80% | 3 | 60% |
| 2 | 3 | 8 | 1 | 85% | 1.5 | 60% |
| 3 | 3 | 9 | 1 | 90% | 1.5 | 60% |
| 4 | 3 | 10 | 1 | 90% | 1.5 | 60% |
| 5 | 3 | 10 | 1 | 90% | 1.5 | 60% |
| 6 | 3 | 10 | 1 | 95% | 1.5 | 60% |
| 7 | 3 | 10 | 1 | 100% | 1.5 | 60% |

Abbreviations: Min, minutes; dur, duration; int, intensity

### Biochemical Assays

Approximately 30 mg of muscle tissue that was flash-frozen in foil was retrieved from - 80°C, weighed using an analytical scale, and homogenized in a sucrose homogenization buffer using a glass Dounce homogenizer according to Spinazzi et al. (22). Samples were centrifuged at 600 × g for 10 minutes at 4°C. Protein concentrations from the resulting supernatants were determined using a commercially available BCA kit (Thermo Fisher Scientific, Waltham, MA, USA). Supernatants were then used for citrate synthase (CS) activity and western blotting.

### Western blotting

Muscle supernatants were prepared for Western blotting using 4x Laemmli buffer and deionized water (diH_2_O) at equal protein concentration. Ten microliters of sample were pipetted onto SDS gels (4%–15% Criterion TGX Stain-free gels; Bio-Rad Laboratories; Hercules, CA, USA), and proteins were separated by electrophoresis (200 V for approximately 40 minutes). Proteins were then transferred to preactivated PVDF membranes (Bio-Rad Laboratories) for 2 hours at 200 mA. Gels were then Ponceau stained for 10 min, washed with diH_2_O for 30 seconds, dried, and digitally imaged (ChemiDoc Touch, Bio-Rad). Following Ponceau imaging, membranes were reactivated in methanol, blocked with nonfat milk for 1 hour, washed three times in Tris-buffered saline with Tween 20 (TBST) and incubated with primary antibodies overnight (1:2000 v/v dilution in TBST with 5% BSA). Primary antibodies were used to detect the following: total OXPHOS rodent (Abcam Cat# ab110413, RRID:AB_2629281), PGC-1α (GeneTex Cat# GTX37356, RRID:AB_11175466), NRF1 (GeneTex Cat# GTX103179, RRID:AB_11168915), TFAM (Abnova Corporation Cat# H00007019-D01P, RRID:AB_1715621), MFN1 (Cell Signaling Technology Cat# 14739, RRID:AB_2744531), MFN2 (BioVision Cat# 3882– 100, RRID:AB_2142625), DRP1 (Novus Cat# NB110-55288SS, RRID:AB_921147), PINK1 (Cell Signaling Technology Cat# 6946, RRID:AB_11179069), and PARKIN (Cell Signaling Technology Cat# 2132, RRID:AB_10693040). Following primary antibody incubations, membranes were washed three times in TBST for 5 minutes and incubated for 1 hour with horseradish peroxidase-conjugated anti-rabbit IgG (Cell Signaling Technology Cat# 7074, RRID:AB_2099233) or anti-mouse IgG (Cell Signaling Technology Cat# 7076, RRID:AB_330924). Membranes were then washed in TBST, developed using chemiluminescent substrate (Millipore; Burlington, MA, USA), and digitally imaged. Raw target band densities were obtained and normalized by Ponceau densitometry values.

### RNA isolation and cDNA synthesis for qPCR analysis

Approximately 10 mg of muscle tissue that was flash-frozen in foil was retrieved from - 80°C, weighed using an analytical scale, homogenized in Ribozol (Ameresco, Solon, OH, USA), and RNA was isolated according to manufacturer’s instructions. RNA concentrations were determined in duplicate using a NanoDrop Lite (Thermo Fisher Scientific, Walthan, MA, USA), and total RNA was determined by normalizing the RNA values to muscle mass homogenized (i.e., µg/mg wet tissue). In an attempt to account for muscle size changes, “absolute” RNA content was estimated by multiplying relative total RNA by mixed fiber cross-sectional area (fCSA) determined by immunohistochemistry (described later) and by vastus lateralis thickness.

For gene expression analyses, 2 µg of cDNA was synthesized using a commercial qScript cDNA SuperMix (Quanta Biosciences, Gaithersburg, MD, USA). RT-qPCR was performed in an RT-PCR thermal cycler (Bio-Rad) using SYBR green-based methods with gene-specific primers designed with primer designer software (Primer3Plus, Cambridge, MA, USA). For all primer sets, pilot qPCR reactions and melt data indicated that only one amplicon was present. The forward and reverse primer sequences of all genes are listed in Table 4. Fold-change values were determined using the 2^ΔΔCq^ method, where 2^ΔCq^ = 2^(housekeeping gene (HKG) Cq - gene of interest Cq) and 2^ΔΔCq^ (or fold change) = (2^ΔCq^ value/2^ΔCq^ average of baseline values). The geometric mean of glyceraldehyde-3-phosphate dehydrogenase (GAPDH) and valosin-containing protein (VCP) was used as the HKG normalizer.

**Table 4.**
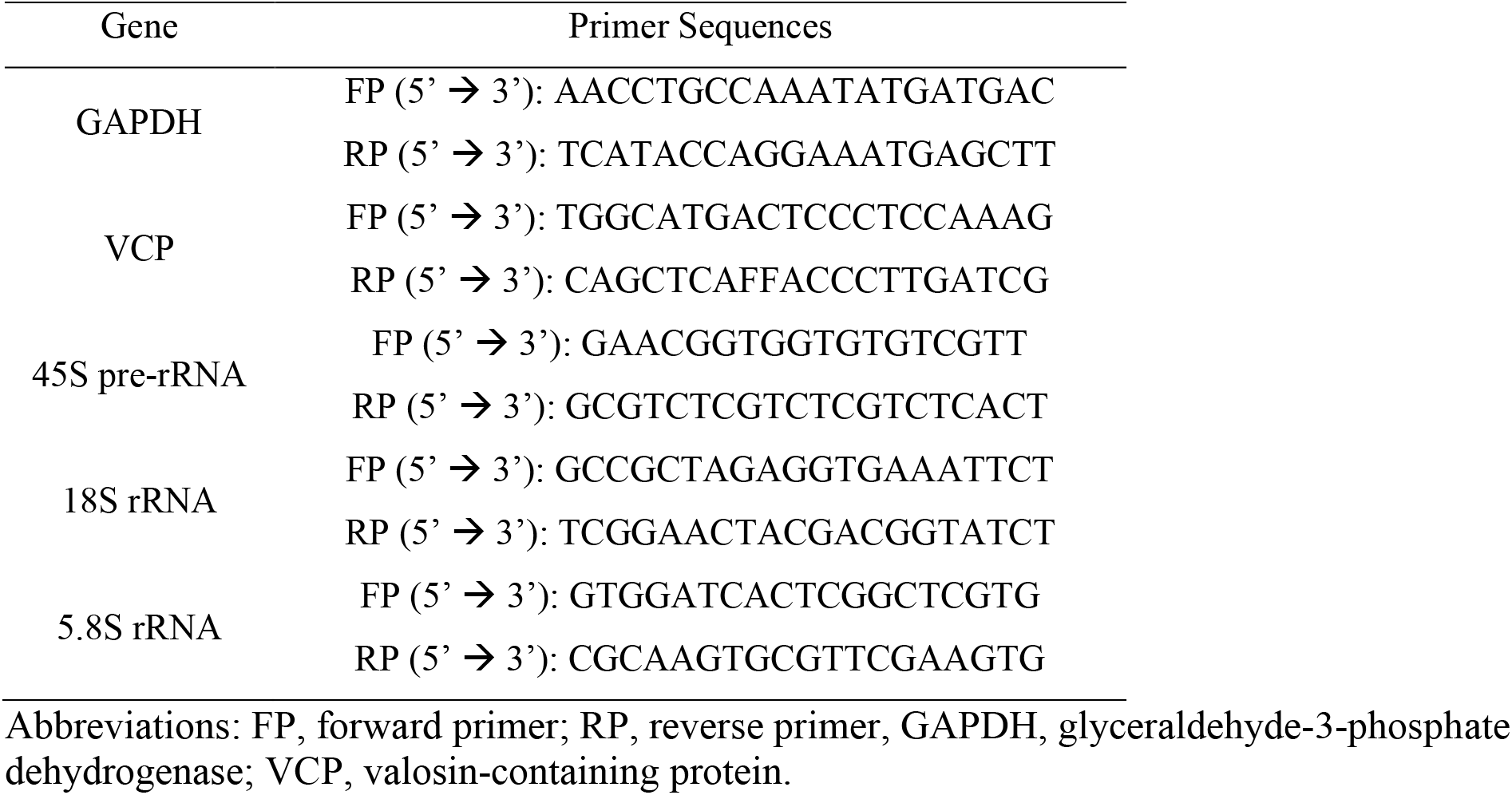
qPCR primer sequences.

### Citrate synthase activity

Citrate synthase activity was determined by monitoring the increase in absorbance at 412 nm from the reduction of 5,5′-dithiobis (2-nitrobenzoic acid) coupled to the reduction of acetyl-CoA (23). Similar to what was listed above in regard to estimating absolute RNA content changes, “absolute” CS activity was calculated by multiplying maximal CS activity (relative) by mixed fCSA and by vastus lateralis thickness.

### Immunohistochemistry (IHC)

A portion of skeletal muscle samples preserved in OCT were sectioned at 7 µm thickness using a cryotome (Leica Biosystems; Buffalo Grove, IL, United States) and adhered to positively charged histology slides. Slides were then stored at -80°C until batch processing. Slides were mounted in a manner that all time-points for each participant were analyzed concomitantly to avoid batch-to-batch variation.

For fiber type-specific fCSA and myonuclei number quantification, slides were air-dried for 90-120 minutes prior to a 5-minute acetone incubation at -20°C. Slides were washed 3×5 minutes in 1x phosphate buffered saline (PBS) and incubated with 3% H_2_O_2_ for 10 minutes at room temperature. After washing, slides were incubated with autofluorescence quenching reagent for 1 min (TrueBlack; Biotium, Fremont, CA, USA). Slides were washed again in PBS and blocked with a 5% goat serum and 2.5% horse serum solution for 1 hour at room temperature. After blocking, slides were incubated overnight at 4°C with a primary antibody cocktail containing 1:20 Mandra (dystrophin) (Developmental Studies Hybridoma Bank; Iowa City, IA, USA) + 1:100 BA-D5 (Myosin Heavy Chain I) (Developmental Studies Hybridoma Bank; Iowa City, IA, USA) + 2.5% horse serum in PBS. The following day, sections were incubated for 1 hour with a secondary antibody cocktail containing 1:250 anti-mouse IgG1 AF594 (Thermo Fisher Scientific; Waltham, MA, USA; cat. no. A-21125) + anti-mouse IgG2b AF488 (Thermo Fisher Scientific; Waltham, MA, USA; cat. no. A-21141) in PBS. Slides were then washed and stained with 1:10,000 DAPI (4=,6-diamidino-2-phenylindole, Thermo Fisher Scientific; catalog #: D3571) for 15 minutes at room temperature before coverslips were applied using PBS + glycerol as mounting medium.

For fiber type-specific satellite cell content quantification, a similar protocol was used. However, additional steps were performed to amplify satellite cells. After blocking slides with 5% goat serum and 2.5% horse serum, slides were blocked with streptavidin and biotin solutions at room temperature for 15 minutes each. Thereafter, slides were incubated overnight at 4°C with primary antibody cocktail containing 1:20 Mandra (dystrophin) (Developmental Studies Hybridoma Bank; Iowa City, IA, USA) + 1:100 BA-D5 (Myosin Heavy Chain I) (Developmental Studies Hybridoma Bank; Iowa City, IA, USA) + 1:20 PAX7 (Developmental Studies Hybridoma Bank; Iowa City, IA, USA) + 2.5% horse serum in PBS. The following day, slides were incubated for 90 minutes in secondary 1:1000 biotin solution (anti-mouse IgG1, Jackson ImmunoResearch; West Grove, PA, USA), followed by a 60-minute incubation with secondary 1:500 streptavidin (SA-HRP, Thermo Fisher Scientific; catalog #: S-911), and a 20-minute incubation with 1:200 tyramide AF555 (Thermo Fisher Scientific, catalog #: B-40957).

For fiber type-specific mitochondrial content, the translocase of outer mitochondrial membrane 20 (TOMM20) protein was stained as previously described and validated (14) using serial sections. The protocol used for mitochondrial staining was similar to those used for fCSA, myonuclei, and satellite cell determination, although the blocking solution included 0.1% Triton X. The primary antibody cocktail included 1:20 Mandy s8 (dystrophin) (Developmental Studies Hybridoma Bank; Iowa City, IA, USA) and 1:200 TOMM20 (Abcam; Cambridge, MA, USA, ab186735) in 5% bovine serum albumin. The following day, slides were incubated for 1 hour with a secondary antibody cocktail: 1:250 anti-rabbit IgG Texas Red 594 (Vector Labs, Newark, CA, USA; TI-1000) + anti-mouse IgG2b AF488 (Thermo Fisher Scientific; Waltham, MA, USA; cat. no. A-21141) in PBS. Slides were then washed in PBS and coverslips were applied using PBS + glycerol as mounting medium.

Single fiber analyses were also performed to quantify myonuclei content. As stated above, muscle tissue (∼10 mg) was fixed in 4% paraformaldehyde for 48 hours at room temperature following biopsies. Tissue was then washed in PBS and stored at 4°C until batch processing. Tissue was subsequently incubated in 40% NaOH in slow rotation for approximately 2 hours to facilitate extracellular matrix digestion and myofiber disaggregation. Tissue was then washed in PBS through a 40 µm cell strainer and transferred to PBS. Small myofiber bundles were mechanically teased apart under a light microscope, placed in PBS, and centrifuged at 13,000 rpm. PBS was removed and myofibers were stained with DAPI for 15 minutes. Individual myofibers were mounted with PBS-glycerol solution on positively charged slides. The number of fibers analyzed were as follows (mean ± SD): RT+ET group, T1: 19±3, T2: 20±1, T3: 20±1; ET-only group, T2: 20±1, T3: 20±1.

Following mounting, digital images for each analysis were captured with a fluorescence microscope (Nikon Instruments) using the 20x objective. Fiber type-specific fCSA and myonuclear number were analyzed using the open-sourced software MyoVision (24). Satellite cells were manually quantified using NIKON NIS Elements software (Nikon Instruments, Melville, NY, USA) and are reported as PAX7 positive per 100 fibers. Mitochondrial content was determined in serial sections using ImageJ (NIH) as previously described (14) and reported as percentage change from baseline. In short, the red channel of TOMM20 images was converted to grayscale and a threshold function was applied. Fibers were then manually traced and mitochondrial area was determined as a percentage of the fiber area. Absolute mitochondrial content via TOMM20 was estimated by multiplying the percentage of TOMM20 by mixed, type I, and type II fiber cross-sectional areas. For single fiber nuclei content, a brightfield image was taken to determine fiber border. Thereafter, three images of the DAPI filter were taken at different depths to capture the maximum number of nuclei. Single fiber and nuclei measurements were made by a blinded investigator using ImageJ (NIH). Myonuclei content is expressed as number of nuclei per 100 µm. Single fiber myonuclear domain (MND) was calculated by dividing the fiber segment volume (µm^3^) by the total number of myonuclei (25).

### Statistical Analysis

Statistics were performed using RStudio Version 2022.12.0. Shapiro-Wilk tests were used to assess the distribution of data for each dependent variable. Two separate analyses were conducted. First, dependent variable responses to resistance training (T1 × T2) in the RT+ET group were analyzed using dependent samples t-tests (for normally distributed data) or Wilcoxon signed rank tests (for non-normally distributed data). Adaptations to endurance training (T2 × T3) in both groups were analyzed using two-way analysis of variance (ANOVA) tests, followed by Tukey post-hoc tests when appropriate. Associations between select variables were also conducted using Pearson’s or Spearman’s correlations. Statistical significance was established at p < 0.05. All data are expressed as mean ± standard deviation (SD) values, and 95% confidence intervals are presented for statistically significant differences.

## RESULTS

### Strength and body composition changes

Participants in the RT+ET group significantly increased 3RM values in leg press (T1: 173 ± 54 kg, T2: 259 ± 56 kg, ± 95% CI [25], p<0.001), bench press (T1: 55 ± 13 kg, T2: 64 ± 14 kg, ± 95% CI [3], p<0.001) and leg extension (T1: 97 ± 25 kg, T2: 128 ± 20 kg, ± 95% CI [8], p<0.001) in response to RT. Further, participants significantly increased lean body mass (+1.4 kg, ± 95% CI [0.9], p=0.005) and VL thickness (+0.22 cm, ± 95% CI [0.08], p<0.001), and decreased body fat percentage (-0.9% ± 95% CI [0.7], p=0.028) in response to RT, whereas no significant changes in body mass (p=0.052) and fat mass (p=0.288) occurred.

In response to ET in both groups, there were no significant effects of group (G), time (T), or interaction (GxT) for body mass (G, p=0.400; T, p=0.111; GxT, p=0.870; Fig. 2A), lean body mass (G, p=0.797; T, p=0.242; GxT, p=0.324; Fig. 2B), or fat mass (G, p=0.118; T, p=0.199; GxT, p=0.818; Fig. 2C). A significant effect of G (p=0.048) and T (p<0.001), but no GxT (p=0.087), was evident for body fat percentage (Fig. 2D). Body fat percentage was higher in the ET-only group (6.4% ± 95% CI [5.9]) and decreased over time (0.9% ± 95% CI [0.5]). No significant main effects of G (p=0.685) or T (p=0.053) were evident for VL thickness (Fig. 2E), but there was a significant GxT (p=0.003). Notably, a decrease in VL thickness occurred in the RT+ET group from T2 to T3 (-0.12 cm ± 95% CI [0.06], p=0.007), but not in the ET group (p=0.805).

**Figure 2.**
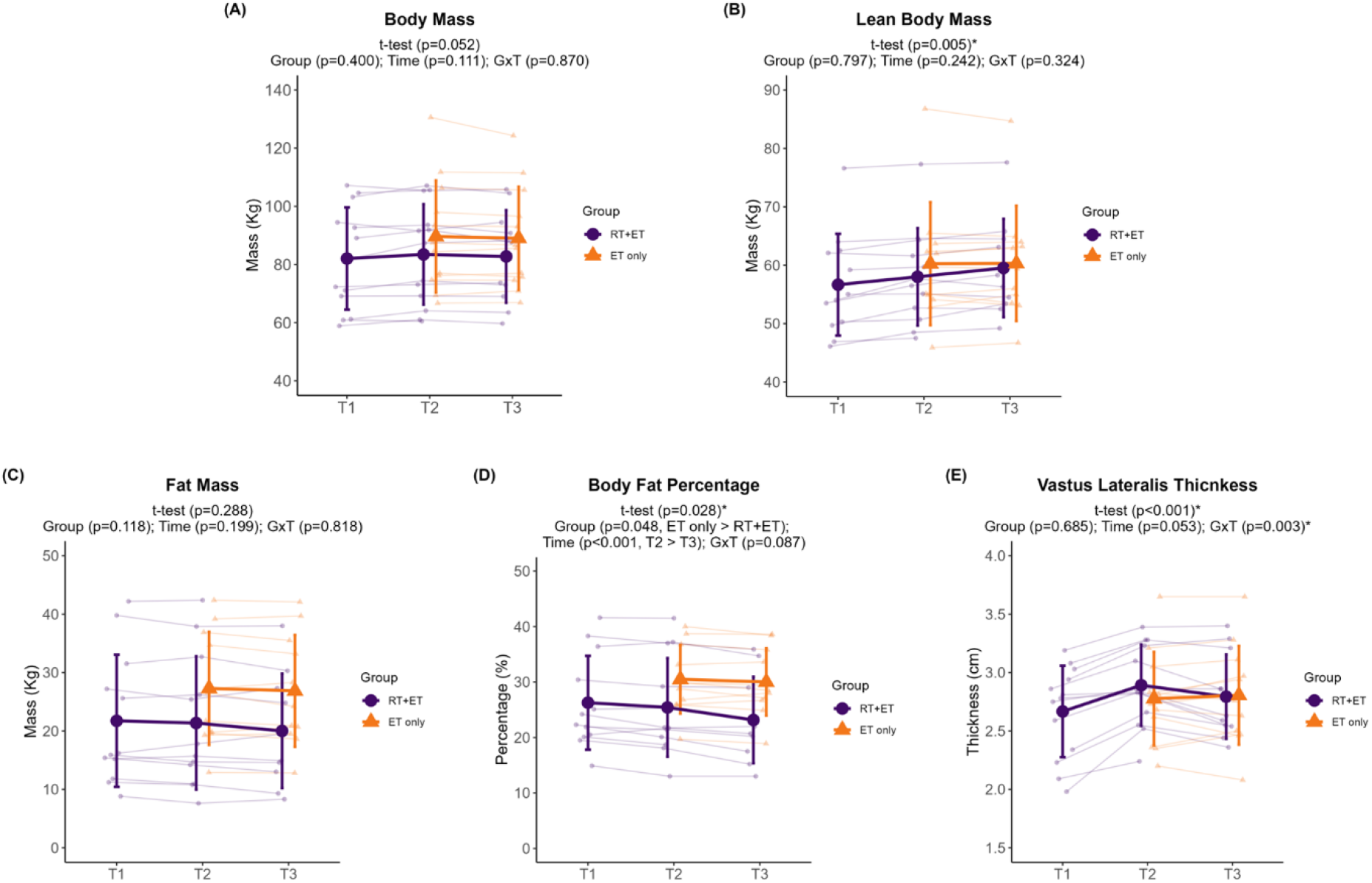
Body composition variables response to RT and ET. (A) Body mass. (B) Lean body mass. (C) Fat mass. (D) Body fat percentage. (E) Vastus lateralis thickness. T1 = Pre-RT; T2 = Pre-ET; T3 = Post-ET. Data are expressed as mean ± SD, and individual respondent values are also depicted. Abbreviations: RT+ET, group that performed 7 weeks of resistance training followed by 7 weeks of endurance training; ET-only, group that performed 7 weeks of endurance training only; GxT, group x time interaction. Notes: t-test p-values are for the RT period in the RT+ET group, and the two-way ANOVA main effect and interaction p-values are for the ET period in both groups.

### Endurance performance

In response to RT, RT+ET participants presented no significant changes in any of the aerobic performance variables (absolute VO_2_max, p=0.058; relative VO_2_max, p=0.794; time to exhaustion, p=0.624; OBLA, p=0.929).

In response to ET in both groups, there was a significant increase over time in all endurance performance variables. Absolute VO_2_max increased 0.38 L/min (± 95% CI [0.07], p<0.001), with no significant main effect of G (p=0.674) or GxT (p=0.154) (Fig. 3A). Relative VO_2_max increased 4.74 ml/kg/min (± 95% CI [0.83], p<0.001), with no significant main effect of G (p=0.108) or GxT (p=0.082) (Fig. 3B). Time to exhaustion during the VO_2_max treadmill test increased 86 seconds (± 95% CI [23], p<0.001), with no significant main effect of G (p=0.430) or GxT (p=0.257) (Fig. 3C). Speed at OBLA increased 0.8 km/h (± 95% CI [0.3], p<0.001), with no significant main effect of G (p=0.354) or GxT (p=0.315) (Fig. 3D).

**Figure 3.**
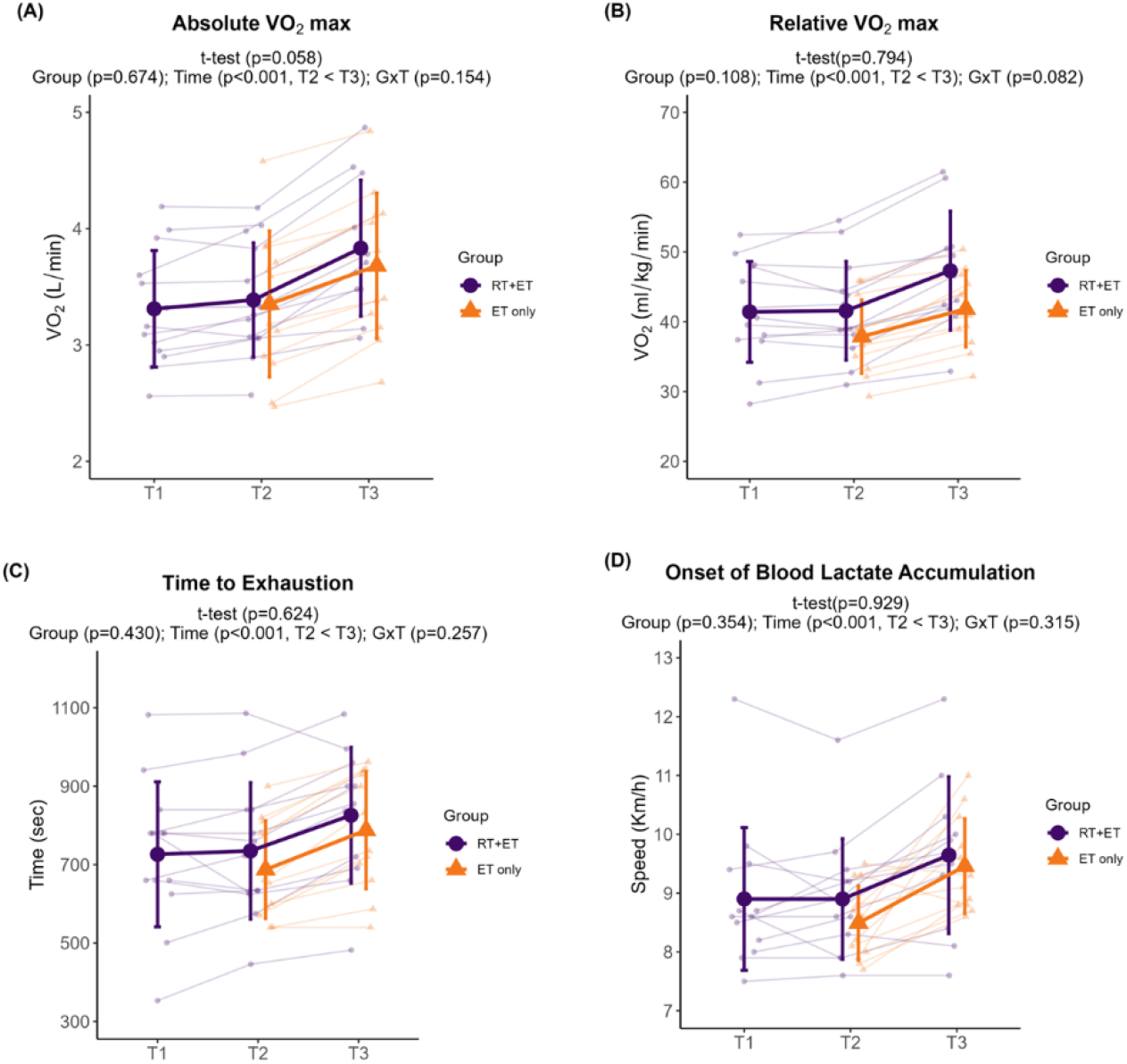
Endurance performance variables response to RT and ET. (A) Absolute VO_2_max. (B) Relative VO_2_max. (C) Time to exhaustion. (D) Onset of blood lactate accumulation. T1 = Pre-RT; T2 = Pre-ET; T3 = Post-ET. Data are expressed as mean ± SD, and individual respondent values are also depicted. Abbreviations: RT+ET, group that performed 7 weeks of resistance training followed by 7 weeks of endurance training; ET-only, group that performed 7 weeks of endurance training only; GxT, group x time interaction. Notes: t-test p-values are for the RT period in the RT+ET group, and the two-way ANOVA main effect and interaction p-values are for the ET period in both groups.

### Mitochondrial remodeling

In response to RT, RT+ET participants exhibited no significant increases in the mitochondrial protein complexes (CI, p=0.659; CII, p=0.110; CIII, p=0.408; CIV, p=0.803; CV, p=0.103). Regarding markers of mitochondrial biogenesis, PGC-1α remained unaltered (p=0.455), NRF-1 increased (0.22 a.u., ± 95% CI [0.10], p<0.001), and TFAM decreased (0.39 a.u., ± 95% CI [0.26], p=0.006). The mitophagy marker PINK1 remained unaltered (p=0.143), while PARKIN significantly decreased (0.36 a.u., ± 95% CI [0.22], p=0.004). Markers of mitochondrial fusion did not change (MFN1, p=0.703; MFN2, p=0.187), but DRP1, a marker of mitochondrial fission, significantly increased (0.19 a.u., ± 95% CI [0.13], p=0.006).

In response to ET in both groups, there were significant increases in mitochondrial protein complexes I-IV from T2 to T3 (CI: 0.23 ± 95% CI [0.15], p=0.006; CII: 0.25 ± 95% CI [0.15], p=0.004; CIII: 0.42 ± 95% CI [0.34], p=0.026; CIV: 0.32 ± 95% CI [0.23], p=0.012) (Fig. 4A). There was no significant main effect of G or GxT for complexes I-III (CI: G, p=0.917 GxT, p=0.062; CII: G, p=0.507, GxT, p=0.177; CIII: G, p=0.987, GxT, p=0.173). For complex IV, there was no significant main effect of G (p=0.878), but a significant GxT (p=0.039), where ET-only was higher at T3 compared to T2 (0.58 a.u., ± 95% CI [0.32], p=0.011). No significant main effects of G (p=0.398), T (p=0.173), or GxT (p=0.164) were evident for complex V. Regarding mitophagy markers (Fig. 4B), no significant main effects of G (p=0.840) or T (p=0.778), or GxT (p=0.140) were evident for PINK1 protein levels. PARKIN levels significantly decreased from T2 to T3 (0.23 a.u. ± 95% CI [0.16], p=0.010), but there was no significant main effect of G (p=0.175) or GxT (p=0.389). Regarding mitochondrial biogenesis markers (Fig. 4C), there were no main effects of G (p=0.214) or T (p=0.926) for PGC-1α, but there was a significant GxT (p=0.001), where ET-only was higher at T3 compared to RT+ET at T3 (0.26 ± 95% CI [0.17], p=0.041). There were no significant main effects of G, T, or GxT for NRF1 (G, p=0.192; T, p=0.912; GxT, p=0.099) or TFAM (G, p=0.062; T, p=0.078; GxT, p=0.458). Regarding mitochondrial dynamics markers (Fig. 4D), no significant main effects of G, T, or GxT were evident for MFN1 (G, p=0.555; T, p=0.079; GxT, p=0.717) or DRP1 (G, p=0.857; T, p=0.122; GxT, p=0.096). MFN2 significantly increased from T2 to T3 (0.03 ± 95% CI [0.02], p=0.030), but there was no significant main effect of G (p=0.066) or GxT (p=0.161).

**Figure 4.**
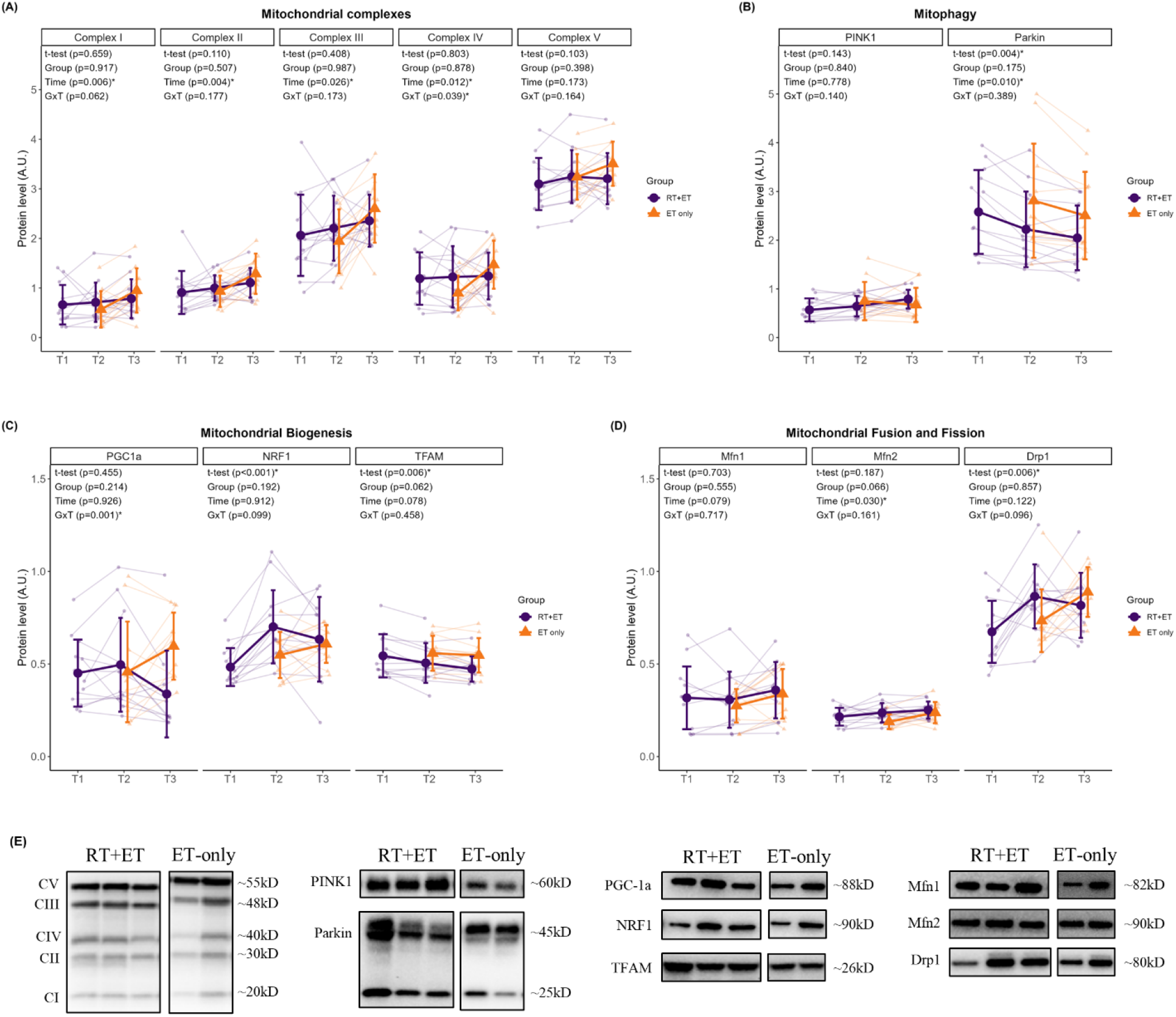
Markers of mitochondrial remodeling response to RT and ET. (A) Mitochondrial complexes. (B) Mitophagy. (C) Mitochondrial biogenesis. (D) Mitochondrial fusion and fission. (E) Representative Western blots. T1 = Pre-RT; T2 = Pre-ET; T3 = Post-ET. Data are expressed as mean ± SD, and individual respondent values are also depicted. Abbreviations: RT+ET, group that performed 7 weeks of resistance training followed by 7 weeks of endurance training; ET-only, group that performed 7 weeks of endurance training only; GxT, group x time interaction. Notes: t-test p-values are for the RT period in the RT+ET group, and the two-way ANOVA main effect and interaction p-values are for the ET period in both groups.

### Ribosome content

In response to RT, there were no significant increases in relative total RNA levels in the RT+ET group (p=0.443). However, when accounting for muscle size, there was a significant increase in absolute RNA estimated by fCSA (944,387 a.u. ± 95% CI [866,900], p=0.035), although there was no increase in absolute RNA estimated by VL thickness (p=0.175). 45S pre-rRNA remained unaltered (p=0.124), while 18S rRNA (0.46 ± 95% CI [0.40], p=0.006) and 5.8S rRNA (0.43 ± 95% CI [0.42], p=0.048) significantly increased with RT.

In response to ET in both groups, there were no significant effects of G or T for relative total RNA (G, p=0.690; T, p=0.779), absolute RNA by fCSA (G, p=0.740; T, p=0.451), or absolute RNA by VL thickness (G, p=0.815; T, p=0.896) (Fig. 5A-C). However, significant GxT were evident for all variables. Relative total RNA was higher in RT+ET group at T2 compared to ET-only group at T2 (181 µg ± 95% CI [118.4], p=0.004), absolute RNA by fCSA post-hoc tests were p>0.050 for all comparisons, and absolute RNA by VL thickness was higher in RT+ET group at T2 compared to ET-only group at T2 (321.4 µg.cm ± 95% CI [321.4], p=0.010). For rRNA transcript levels (Fig. 5D), there were no significant main effects of G or T, and no significant GxT (45S pre-rRNA: G, p=0.794; T, p=0.461; GxT, p=0.873; 18S rRNA: G, p=0.501; T, p=0.661; GxT, p=0.292; 5.8S rRNA: G, p=0.223; T, p=0.485; GxT, p=0.095).

**Figure 5.**
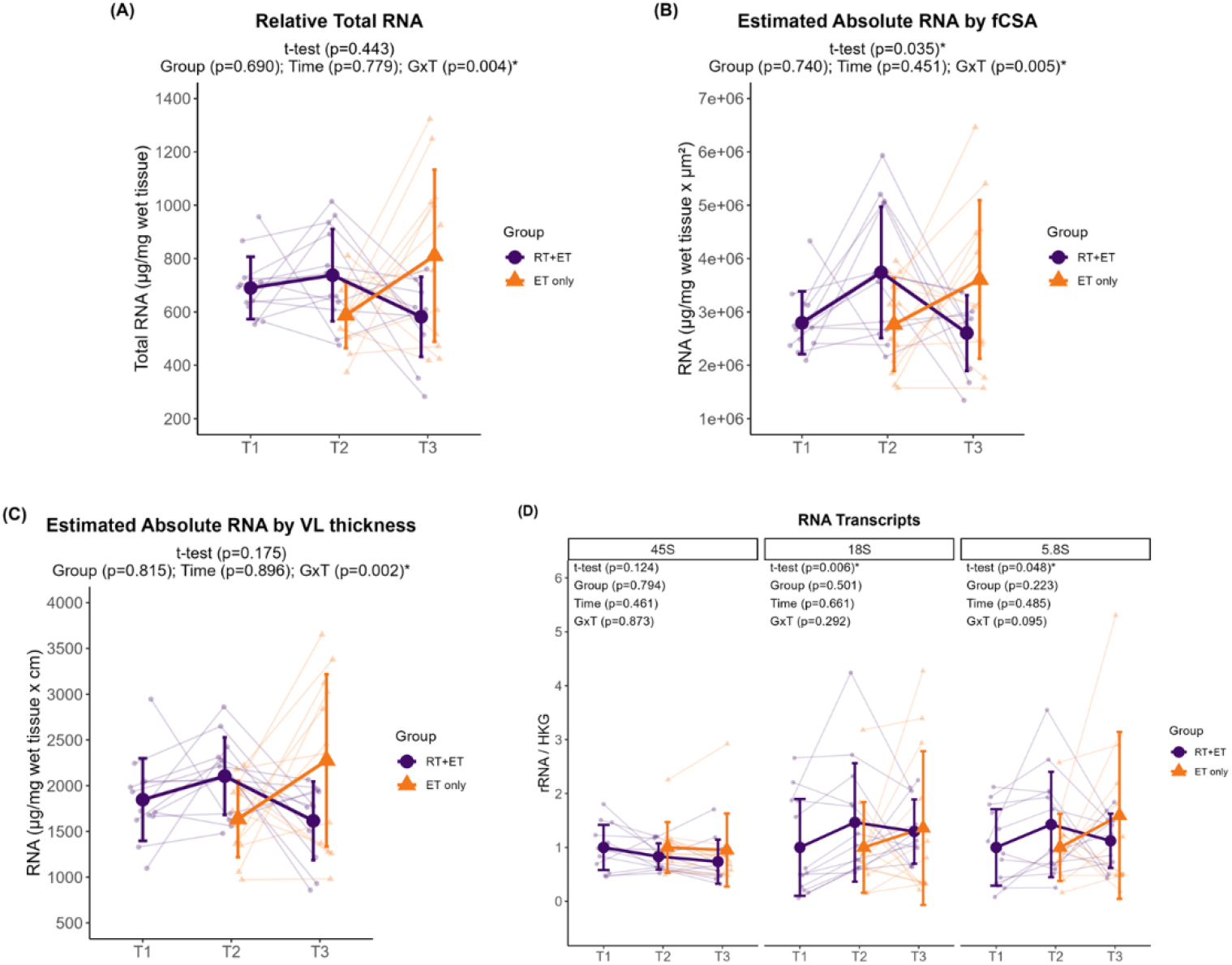
Markers of ribosome content response to RT and ET. (A) Total RNA concentrations. (B) Estimated absolute RNA content (adjusted for mixed fiber cross-sectional area values). (C) Estimated absolute RNA content (adjusted for VL thickness values) (D) Ribosomal RNA transcripts. T1 = Pre-RT; T2 = Pre-ET; T3 = Post-ET. Data are expressed as mean ± SD, and individual respondent values are also depicted. Abbreviations: RT+ET, group that performed 7 weeks of resistance training followed by 7 weeks of endurance training; ET-only, group that performed 7 weeks of endurance training only; GxT, group x time interaction. Notes: t-test p-values are for the RT period in the RT+ET group, and the two-way ANOVA main effect and interaction p-values are for the ET period in both groups.

### Immunohistochemistry

#### Fiber cross-sectional area

In response to RT, there were significant increases in mixed (757 µm^2^ ± 95% CI [455], p=0.002) and type II (949 µm^2^ ± 95% CI [527] p<0.001) fCSA in the RT+ET group. However, there were no significant changes in type I fCSA (p=0.129). In response to ET in both groups, there were no significant main effects of G, T, or GxT in mixed (G, p=0.772; T, p=0.087; GxT, p=0.413), type I (G, p=0.668; T, p=0.396; GxT, p=0.992), or type II (G, p=0.497; T, p=0.088; GxT, p=0.340) fCSA (Fig. 6A).

**Figure 6.**
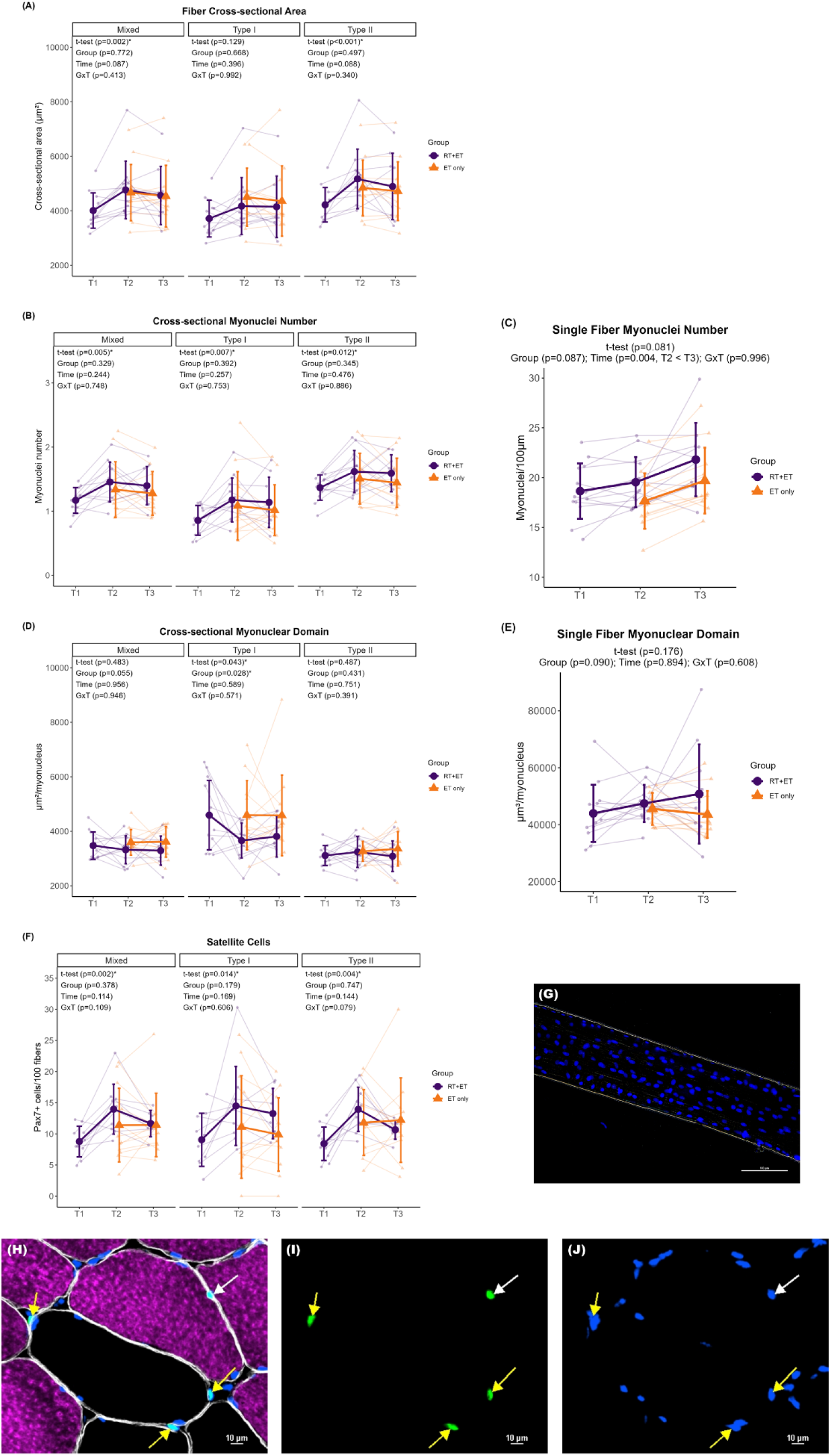
Fiber cross-sectional area, myonuclei and satellite cell number, and myonuclear domain responses to RT and ET. (A) Fiber cross-sectional area. (B) Cross-sectional myonuclei number. (C) Single fiber myonuclei number. (D) Cross-sectional myonuclear domain. (E) Single fiber myonuclear domain. (F) Satellite cells content. (G) Single fiber representative image. (H-J) Representative images of cross-sectional staining. (H) Dystrophin (white), MHCI (magenta), DAPI (blue), Pax7 (green). (I) Pax7. (J) Pax7 + DAPI. T1 = Pre-RT; T2 = Pre-ET; T3 = Post-ET. Data are expressed as mean ± SD, and individual respondent values are also depicted. Abbreviations: RT+ET, group that performed 7 weeks of resistance training followed by 7 weeks of endurance training; ET-only, group that performed 7 weeks of endurance training only; GxT, group x time interaction. Notes: t-test p-values are for the RT period in the RT+ET group, and the two-way ANOVA main effect and interaction p-values are for the ET period in both groups.

#### Myonuclear number

In response to RT, myonuclear number in cross-section increased in mixed (0.3 ± 95% CI [0.2], p=0.005), type I (0.3 ± 95% CI [0.2], p=0.007), and type II (0.3 ± 95% CI [0.2], p=0.012) in the RT+ET group. Myonuclear number as quantified through single fiber analysis, however, did not reach statistical significance (p=0.081). In response to ET in both groups, there were no significant main effects of G, T, or GxT for mixed (G, p=0.329; T, p=0.244; GxT, p=0.748), type I (G, p=0.392; T, p=0.257; GxT, p=0.753), or type II (G, p=0.345; T, 0.476; GxT, p=0.886) myonuclear number in cross-section (Fig. 6B). Single fiber myonuclei number increased from T2 to T3 (2.0 ± 95% CI [1.2], p=0.004), with no main effect of G (p=0.087) or GxT (p=0.996) (Fig. 6C).

#### Myonuclear domain (MND)

In response to RT, there was a decrease in cross-sectional MND in type I fibers (934 µm^2^/myonucleus ± 95% CI [900], p=0.043) in the RT+ET group, but not in mixed (p=0.483) or type II fibers (p=0.487). However, MND values assessed through single fiber analysis exhibited no significant change (p=0.176). In response to ET in both groups, ET-only exhibited greater cross-sectional MND values in type I fibers compared to RT+ET group (G effect: 941 µm^2^/myonucleus ± 95% CI [741], p=0.028). There was no main effect of T (p=0.589) or GxT (p=0.571) for type I MND (Fig. 6D). In addition, there were no significant main effects of G, or T, or GxT in mixed (G, p=0.055; T, p=0.956; GxT, p=0.946) or type II (G, p=0.431; T, p=0.751; GxT, p=0.391) fibers. For single fiber MND, there were no significant effects of G (p=0.090), T (p=0.894), or GxT (p=0.608) (Fig. 6E).

#### Satellite Cells

In response to RT, there was a significant increase in satellite cell content in mixed (5.2 ± 95% CI [2.7], p=0.002), type I (5.4 ± 95% CI [4.9], p=0.014), and type II (5.5 ± 95% CI [3.1], p=0.004) fibers in the RT+ET group. In response to ET in both groups, there were no significant main effects of G or T, or GxT for mixed (G, p=0.378; T, p=0.114; GxT, p=0.109), type I (G, p=0.179; T, p=0.169; GxT, p=0.606), or type II (G, p=0.747; T, p=0.144; GxT, p=0.079) fibers (Fig. 6F).

### Mitochondrial content assessed using CS activity and TOMM20 IHC

In response to RT, relative CS activity remained unaltered (p=0.152), but absolute CS activity significantly estimated by fCSA (481,561 nmol/min/mg.µm^2^ ± 95% CI [335,161], p=0.009) and by VL thickness (152 nmol/min/mg.cm ± 95% CI [117]) increased in response to RT. There were no significant changes in relative mitochondrial content as assessed through TOMM20 IHC in mixed (p=0.296), type I (p=0.982), or type II (p=0.070) fibers in the RT+ET group. Regarding the estimation of total mitochondrial content, which accounts for fCSA changes, there was a significant increase in type II fiber mitochondrial content (45% ± 95% CI [38], p=0.026) but not in mixed (p=0.113) or type I (p=0.476) fibers.

In response to ET in both groups, mitochondrial content assessed through relative CS activity was higher in the ET-only group compared to RT+ET group (main effect of G: 92.7 mmol/min/mg ± 95% CI [81.3], p=0.036) (Fig. 7A). However, there was no significant main effect of T (p=0.461) or GxT (p=0.162). Additionally, there were no significant effects of G, T, or GxT in total mitochondrial content estimated by mixed fCSA (G, p=0.170; T, p=0.117; GxT, p=0.132) or by VL thickness (G, p=0.074; T, p=0.257; GxT, p=0.080) in response to ET (Fig. 7B-C).

**Figure 7.**
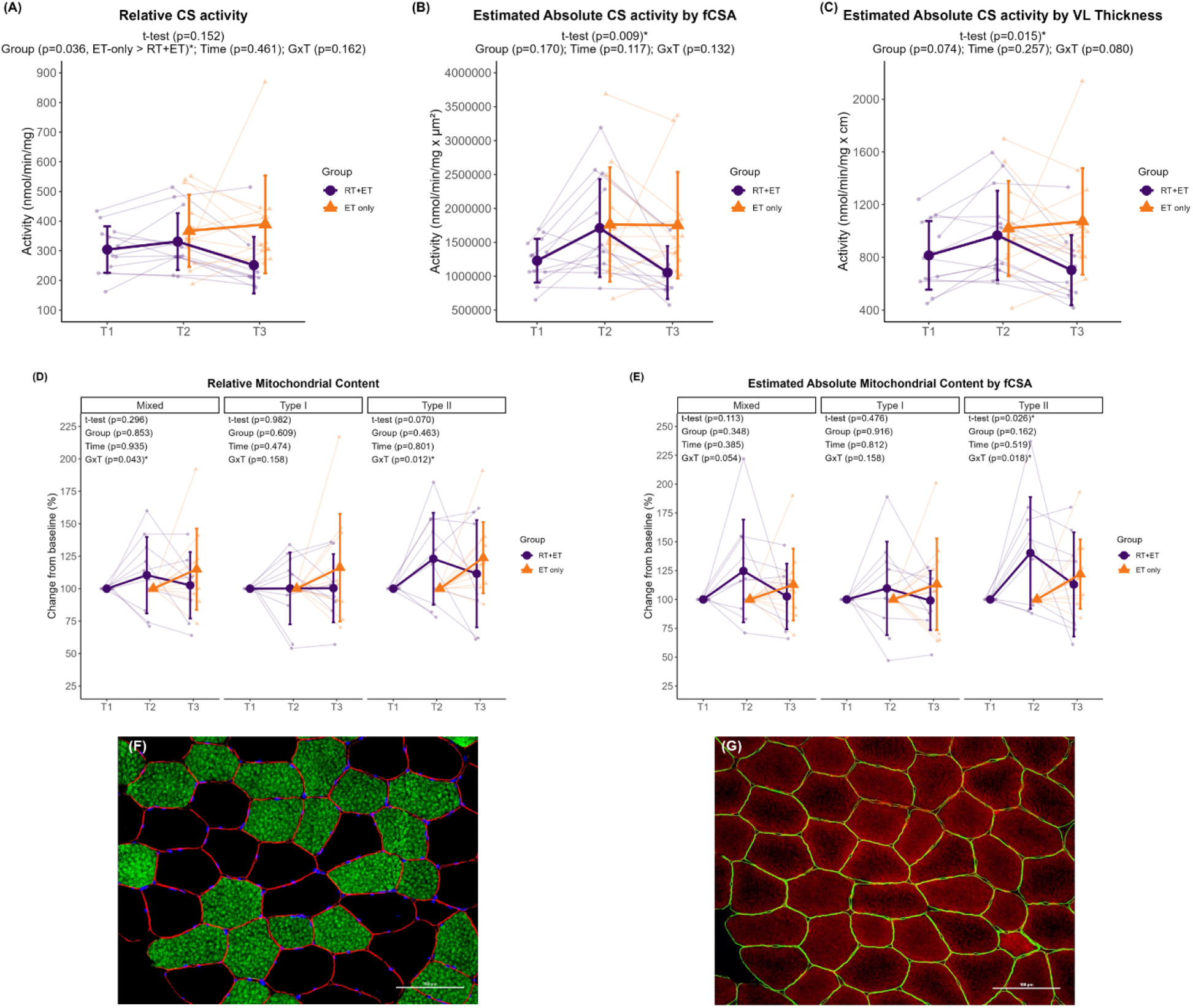
Mitochondrial content responses to RT and ET. (A) Relative maximal CS activity. (B) Total mitochondrial content estimation (via maximal CS activity and mixed fCSA values). (C) Total mitochondrial content estimation (via maximal CS activity and VL thickness). (D) Relative mitochondrial content (via TOMM20 IHC). (E) Total mitochondrial content estimation (via TOMM20 IHC and mixed fCSA values). (F-G) Representative images of serial cross-sectional staining. (F) Dystrophin (red), MHCI (green), DAPI (blue). (G) Dystrophin (green), TOMM20 (red). T1 = Pre-RT; T2 = Pre-ET; T3 = Post-ET. Data are expressed as mean ± SD, and individual respondent values are also depicted. Abbreviations: RT+ET, group that performed 7 weeks of resistance training followed by 7 weeks of endurance training; ET-only, group that performed 7 weeks of endurance training only; GxT, group x time interaction. Notes: t-test p-values are for the RT period in the RT+ET group, and the two-way ANOVA main effect and interaction p-values are for the ET period in both groups.

There were significant GxT for mitochondrial content as assessed through TOMM20 IHC in mixed (p=0.043) and type II fibers (p=0.012). Post-hoc tests for mixed fibers returned p-values > 0.050 for all comparisons. Mitochondrial content in type II fibers was greater in the RT+ET group at T2 compared to ET-only group at T2 (33.8% ± 95% CI [20.2], p=0.036), but there were no significant main effects (for mixed: G, p=0.853 and T, p=0.935; for type II G p=0.463 and T, p=0.801) (Fig. 7D). Type I fibers exhibited no significant main effects of G (p=0.609) or T (p=0.474), or GxT (p=0.158). Similar responses were observed for total mitochondrial content estimations (Fig. 7E). A significant GxT was observed in type II fibers (p=0.018), where mitochondrial content was higher in the RT+ET group at T2 compared to ET-only group at T2 (52.2% ± 95% CI [30.6], p=0.032). No significant main effects of G (p=0.162) or T (p=0.519) were detected for type II fibers. In addition, no significant effects of G, T, or GxT were found in mixed (G, p=0.348; T, p=0.385; GxT, p=0.054) or type I (G, p=0.916; T, p=0.812; GxT, p=0.158).

### Correlations

Correlation between the values at T2 from select variables (e.g., relative total RNA, fCSA, nuclei, SCs) and the percent change of mitochondrial content variables (i.e., relative CS activity, mixed fibers, type I and type II mitochondrial content (TOMM20)) in response to ET were analyzed in a group-specific manner due to the distinctive responses of each group to ET. The only significant correlation found was between relative CS activity and mixed fCSA in the ET-only group (r = 0.64, p=0.028). The correlation between all other variables can be found in Supplementary Figure 1.

## DISCUSSION

Resistance training has long been appreciated for increasing muscle mass and strength, and emerging evidence highlights that RT may also promote positive mitochondrial adaptations. Most studies investigating the differences and interplay between RT and ET adaptations have compared concurrent training to single-mode training, using various experimental designs. To the best of our knowledge, this is the first study to investigate the effects of performing a period of RT-only on the adaptations to a subsequent period of ET-only. Our main findings demonstrate that RT performed prior to ET had no additional benefits to ET adaptations. Moreover, even though both groups improved endurance performance similarly, prior RT seemed to impair most mitochondrial adaptations to subsequent ET.

In the current study, seven weeks of RT elicited adaptations commonly reported in the literature, which demonstrates the effectiveness of the RT protocol adopted herein. Participants in the RT+ET group improved body composition and strength, and increased VL thickness, mixed and type II fCSA, myonuclear number, markers of ribosome content, and satellite cell number. Various methods can be implemented to increase endurance performance, with the most common being MICT and HIIT. High-intensity interval training protocols have been shown to improve endurance performance in as little as two weeks (26). VO_2_max and the speed at lactate threshold are considered key determinants of endurance performance (1, 2). In the current study, relative VO_2_max improved 13.4% in the RT+ET group and 10.6% in the ET-only, which is within the range reported in the literature (6, 27). In addition, the speed at OBLA increased 7% in the RT+ET group and 12% in ET-only group. However, performing a block of RT before initiating ET did not significantly enhance these adaptations to ET. Additionally, much of our molecular data suggest that seven weeks of RT performed prior to seven weeks of ET may interfere with mitochondrial adaptations, and this will be the crux of the remainder of the discussion.

The mitochondrial adaptations to RT are not well defined based on prior literature. For instance, while it is commonly believed that RT is not an effective method to achieve positive mitochondrial adaptations, different researchers have reported increases in markers of mitochondrial content and function in younger (11, 12, 28) and older (13, 29-31) individuals. In the current study, there were no significant changes in mitochondrial protein complex concentrations or markers of mitochondrial content (TOMM20 and CS activity) with RT. However, type II fiber total mitochondrial content (as estimated by considering changes in fCSA) increased. This increase in total, but not relative mitochondrial content, suggests that the expansion of the mitochondrial network occurred in line with type II myofiber size increases. Alternatively stated, we speculate that the metabolic demands of RT did not facilitate mitochondrial expansion per se, but that the expansion of the mitochondrial network occurred in proportion to myofiber size to optimize a mitochondrial-to-myofiber volume ratio.

Satellite cells and myonuclear accretion have been extensively studied in the context of RT and skeletal muscle hypertrophy. Whether or not these events are required for hypertrophy is still a topic of debate (32, 33), albeit satellite cells and myonuclei content are commonly reported to increase with RT (34-36). However, the effects of ET on satellite cells and myonuclear counts have received less attention in the literature. In line with our expectations, seven weeks of RT in the RT+ET group increased mixed, type I and type II myofiber myonuclear number and satellite cell number. However, in both RT+ET and ET-only groups, HIIT training did not elicit significant changes in fCSA, cross-sectional myonuclei or satellite cells number. Our results agree with previous studies that showed no changes in type I and II satellite cell or myonuclear number after different forms of ET (37, 38), and continue to support that RT (but not ET) acts as a stimulus to affect these variables. As with the sparse research examining how ET affects satellite cell number, studies that have examined the effects of ET on ribosome biogenesis markers are also limited. There is a common dichotomous viewpoint whereby RT promotes ribosome biogenesis and ET promotes mitochondrial biogenesis, with an interference effect between the two processes if RT and ET are performed concurrently (39, 40). However, it is possible that untrained individuals can present a generic response to exercise training whereby ribosome and mitochondrial biogenesis can occur in response to both RT and ET (39). In response to the HIIT period in the current study, the RT+ET group presented decreases in ribosome content while the ET-only group presented paradoxical increases in these variables. While these events are difficult to reconcile, the RT+ET response may be related to the cessation of RT and not a response to ET per se, as ribosome content has been previously shown to decrease rapidly upon RT cessation. Hammarström et al. (41), for example, showed a similar decrease in total RNA concentrations (19.3%) after eight days of detraining in humans. Furthermore, Figueiredo et al. (42) found that the decrease in ribosome content during muscle disuse was correlated with the decrease in muscle CSA. Therefore, it is possible that ET did not provide sufficient stimulus for ribosome maintenance, as has been shown by Romero et al. (43) when providing treadmill ET in rats over a 12-week period. Indeed, this hypothesis is speculative given that we do not have time coursed biopsies to examine markers of ribosome degradation in the RT+ET group, and more research is needed in this regard. The increase in ribosome content in the ET-only group is novel and equally as intriguing. Prior rodent work from our laboratory suggests that 12 weeks of HIIT-style treadmill ET increases ribosome biogenesis markers in lieu of decreasing skeletal muscle ribosome content (43). Subsequent work from Figueiredo and collaborators (42) indicated that a bout of resistance exercise upregulates several markers in skeletal muscle indicative of increased ribosome biogenesis, whereas this does not occur in response to a steady-state bout of cycling. Others have also shown that weeks of concurrent training enhances ribosome biogenesis relative to resistance training alone (44). Hence, these prior and our current data suggest that the mode of exercise (e.g., HIIT versus steady state) and (perhaps) species differences may affect the ribosome biogenesis response to ET.

The majority of studies investigating molecular adaptations in response to ET have focused on mitochondrial variables due to their importance in oxidative metabolism. Several studies have shown increased mitochondrial content and function in response to various forms of ET (5, 45, 46). Considering that approximately 98% of the proteins that make up mitochondria are encoded by the nuclear genome (47), we hypothesized that RT-mediated increases in myonuclei and ribosomes would increase both the transcriptional and translational capacity of myofibers, allowing for enhanced mitochondrial adaptations. In fact, Lee and collaborators (15) reported that prior RT facilitated mitochondrial adaptations to a subsequent block of RT in rats. Using both rodent and cell models, these authors also demonstrated that higher myonuclear number was related to a greater expression of mitochondrial genes and proteins in response to exercise. However, even though RT led to increased myonuclei and ribosome content in the current study, most mitochondrial adaptations to subsequent ET were blunted. For example, the protein levels of mitochondrial complexes I-IV in the ET-only group showed increases from 32% to 66%, while the RT+ET group only increased from 1% to 11%. Moreover, mixed fiber relative mitochondrial content increased 15% in the ET-only group but decreased 13% in the RT+ET group. Once more, the reasons for such distinctive responses to ET are difficult to reconcile. However, given that the RT+ET group also exhibited decreases in several other variables (e.g., VL thickness, fCSA, and RNA levels), we speculate that these participants existed in an enhanced catabolic/proteolytic state during the duration of the seven week ET period. In support of this hypothesis are certain lines of evidence that have used stable isotopes to ascertain mixed and fractional synthetic protein turnover rates. It is well-known that resistance exercise acutely stimulates increases in both muscle protein synthesis and breakdown, albeit with chronic training, increases in muscle protein synthesis generally exceed increases in muscle protein breakdown (48-50). These events promote a longer-term net positive in protein balance and result in myofiber and whole-tissue skeletal muscle hypertrophy. On the other hand, while ET increases muscle protein synthesis and breakdown (49, 51), the increases in muscle protein synthesis may be specific to mitochondrial (rather than myofibrillar) protein synthesis (52). In addition, ET has been shown to increase several proteolytic markers in skeletal muscle (53, 54). When considering these prior data and our current observations, it remains possible that the transition from RT to ET in the RT+ET group promoted a sustained elevation in muscle protein breakdown mechanisms while diminishing the protein synthetic response. An ultimate consequence of this shift may have included myofiber atrophy accompanied by a decrease in cellular mitochondrial and ribosome content. While this is an attractive hypothesis to explain several of the RT+ET observations herein, it is speculative and further investigation is needed to confirm this hypothesis.

### Experimental considerations

There are limitations to the current study. First, the n-sizes and biopsy time points were limited in scope. Moreover, only younger adult men were examined herein. Hence, these data should be viewed with these limitations in mind. Additionally, it is important to note that we did not ascertain muscle protein synthesis or breakdown rates, and proteolytic markers were not assayed. As such, much of our speculations regarding the RT+ET adaptations require further inquiry. Markers of mitochondrial function were also not measured in the current study, and it is possible that mitochondrial function improved in response to RT and/or ET. Finally, the inclusion of a control group with a detraining period after RT is lacking, and the inclusion of such a group would have helped distinguish the effects of RT cessation from ET adaptations.

### Conclusions

In conclusion, the results of the present study showed that prior RT had no additional benefits on performance adaptations to ET. Additionally, several mitochondrial adaptations to ET (as well as other molecular outcomes) were blunted in the RT+ET group following the ET period. Whether these maladaptive responses at the molecular level have longer-term functional consequences remains to be determined.

## ACKNOWLEDGEMENTS

Participant compensation costs and most reagent costs were funded by a grant provided by the National Strength and Conditioning Association Foundation to P.H.C.M. M.C.M. was fully supported through a T32 NIH grant (T32GM141739). C.A.L. was supported by the São Paulo Research Foundation 393 (n° 2020/13613-4) and Technological Development (n° 302801/2018-9). Assay costs not covered by the National Strength and Conditioning Foundation were provided through discretionary laboratory funds from M.D.R and A.N.K., and the Auburn School of Kinesiology paid for publishing fees. None of the authors have financial or other conflicts of interest to report regarding these data.

## AUTHOR CONTRIBUTIONS

P.H.C.M. primarily drafted the manuscript and constructed figures. All co-authors were involved in critical aspects of the study regarding data collection and analyses. M.D.R. and A.N.K. provided critical assistance in manuscript preparation. All co-authors edited the manuscript, and all authors approved the final submitted version.

## DATA AVAILABILITY STATEMENT

Several raw data files can be obtained upon reasonable request by emailing the latter co-corresponding/senior author.

